# The information theory formalism unifies the detection of the patterns of sexual selection and assortative mating for both discrete and quantitative traits

**DOI:** 10.1101/2023.08.14.552693

**Authors:** A. Carvajal-Rodríguez

**Author notes:** A. Carvajal-Rodríguez. Departamento de Bioquímica, Genética e Inmunología. Universidade de Vigo, 36310 Vigo, Spain.

## Abstract

Sexual selection plays a crucial role in modern evolutionary theory, offering valuable insight into evolutionary patterns and species diversity. Recently, a comprehensive definition of sexual selection has been proposed, defining it as any selection that arises from fitness differences associated with nonrandom success in the competition for access to gametes for fertilization. Previous research on discrete traits demonstrated that non-random mating can be effectively quantified using Jeffreys (or symmetrized Kullback-Leibler) divergence, capturing information acquired through mating influenced by mutual mating propensities instead of random occurrences. This novel theoretical framework allows for detecting and assessing the strength of sexual selection and assortative mating.

In this study, we aim to achieve two primary objectives. Firstly, we demonstrate the seamless alignment of the previous theoretical development, rooted in information theory and mutual mating propensity, with the aforementioned definition of sexual selection. Secondly, we extend the theory to encompass quantitative traits. Our findings reveal that sexual selection and assortative mating can be quantified effectively for quantitative traits by measuring the information gain relative to the random mating pattern. The connection of the information indices of sexual selection with the classical measures of sexual selection is established.

Additionally, if mating traits are normally distributed, the measure capturing the underlying information of assortative mating is a function of the square of the correlation coefficient, taking values within the non-negative real number set [0, +∞).

It is worth noting that the same divergence measure captures information acquired through mating for both discrete and quantitative traits. This is interesting as it provides a common context and can help simplify the study of sexual selection patterns.

## Introduction

Sexual selection is a fundamental concept in modern evolutionary theory, offering valuable insights into various evolutionary patterns and the diversity of species.

Originally proposed by Darwin in (1871), sexual selection describes the competition among individuals of one sex to secure mates from the other sex. However, the concept of sexual selection has been a subject of controversy since its inception (Alonzo and Servedio, 2019; Andersson, 1994; Lehtonen, 2022; Parker, 2014; Parker and Pizzari, 2015; Prum, 2012; Roughgarden et al., 2015; Vries and Lehtonen, 2023). There is ongoing debate surrounding its precise definition (Fitze et al., 2011), and its role as a key component of modern evolutionary biology has been challenged (Roughgarden et al., 2006; Parker and Pizzari, 2015; but see Shuker, 2010).

Recently, Shuker and Kvarnemo (2021) put forth a comprehensive definition of sexual selection, characterizing it as “any selection that arises from fitness differences associated with nonrandom success in the competition for access to gametes for fertilization”. In essence, a trait falls under sexual selection when variations in that trait are nonrandomly associated with variations in access to gametes, which serve as a limiting resource. From an evolutionary perspective, this process generates a discernible pattern of change in gene frequency, influencing the frequencies of corresponding genotypes and phenotypes. It is important to emphasize the use of the term “pattern of change” when discussing sexual selection, as it helps to clarify whether we are referring to the dynamic process or to the recognizable pattern in genetic or phenotypic changes.

The pattern of sexual selection is caused mainly by two biological mechanisms: mate competition and mate choice (Arnold and Wade, 1984; Endler, 1986; Lewontin et al., 1968; Rolán-Alvarez et al., 2015; Rolán-Alvarez and Caballero, 2000).

The process of mate competition refers, in a broad sense, to access to mating through courtship, intrasexual aggression, competition for limited reproductive resources and post-copulatory mechanisms as sperm competition (Andersson, 1994; Kokko et al., 2012; Wacker and Amundsen, 2014). Mate competition may generate a pattern of sexual selection (a change in frequencies of the trait under study) in the sex that competes.

The process of mate choice refers to the effects of certain traits expressed in one sex that leads to the non-random allocation of reproductive effort by members of the opposite sex (Edward, 2015; Rosenthal, 2017). Mate choice may be mediated by phenotypic (sensorial or behavioral) properties that affect the propensity of individuals to mate with certain phenotypes (Jennions and Petrie, 1997). The observed pattern driven by mate choice, including cryptic female choice, can be a change in trait frequency in the other sex (sexual selection) and/or assortative mating where similar phenotypes or genotypes mate with one another more or less frequently than would be expected under a random mating pattern i.e., it produces a deviation from random mating in mated individuals (Carvajal-Rodríguez, 2018a; Rolán-Alvarez and Caballero, 2000). When the observed mating pattern between similar phenotypes is more frequent than expected by chance, we refer to positive assortative mating and when it is less frequent we refer to negative assortative mating. Sometimes in the literature, positive assortative mating is just referred to as assortative mating and negative assortative mating as disassortative mating. Mate choice when it produces positive assortative mating can have important evolutionary effects as it alters the variance of quantitative traits (Fisher, 1918), can generate sexual selection (Carvajal-Rodríguez, 2018a; de Cara et al., 2008) and under certain conditions, may facilitate local adaptation and speciation even in the presence of gene flow (Cotto and Servedio, 2017; Coyne and Orr, 2004; Fernández-Meirama et al., 2022; Gavrilets, 2004; Kirkpatrick and Ravigné, 2002; Rettelbach et al., 2013; Sachdeva and Barton, 2017; Yukilevich, 2023).

When examining phenotypes involving discrete traits, sexual selection can be identified through mating data using the pair sexual selection statistic (PSS), similarly, assortative mating can be identified by the pair sexual isolation statistic (PSI, Carvajal-Rodríguez and Rolan-Alvarez, 2006; Rolán-Alvarez and Caballero, 2000).

For quantitative traits, almost all measures of sexual selection currently used require estimates of variance in mating or reproductive success (reviewed in Henshaw et al., 2018, 2016; Jones, 2009). In this work I focus on mating and fertilization success, rather than reproductive success (Shuker and Kvarnemo, 2021). On the other side, assortative mating can be assessed by computing the Pearson correlation coefficient within mated pairs (Jiang et al., 2013). However, it is important to note that this measure comes with known problems and limitations (Clancey et al., 2022; Fernández-Meirama et al., 2022).

In a previous study (Carvajal-Rodríguez, 2018a), it was demonstrated that non-random mating can be effectively quantified using the Jeffreys (Kullback-Leibler symmetrized) divergence. This measure captures the information acquired when mating is influenced by mutual mating propensities, rather than occurring randomly. By employing this theoretical framework, it becomes possible to detect and assess the strength of sexual selection and assortative mating (Carvajal-Rodríguez, 2020; Estévez et al., 2020; Lau et al., 2021).

The use of information theory to describe evolutionary processes is not new as it has a long tradition in evolutionary ecology, (see for example Sherwin, 2018; Sherwin et al., 2017; Sherwin and Prat i Fornells, 2019) and population genetics (Frank, 2012; Hledík et al., 2022; Kimura, 1961). Previous work on information theory and non-random mating was done on discrete traits (Carvajal-Rodríguez, 2018a, 2019, 2020). Here we extend that work to quantitative traits to describe patterns of sexual selection and assortative mating in terms of information.

The objective of this work is twofold. Firstly, we aim to demonstrate that the previous theoretical development, which is rooted in information theory and the concept of mutual mating propensity (Carvajal-Rodríguez, 2018a, 2020), seamlessly aligns with the sexual selection definition put forth by Shuker and Kvarnemo (2021). In this context, sexual selection emerges as a consequence of variations in the mutual mating propensity, without requiring specific assumptions about the genetic basis of the involved traits. Secondly, we aim to illustrate that, for quantitative traits, sexual selection is also measured by Jeffreys divergence, which in turn can also be connected to classical metrics such as the opportunity for sexual selection and the standardized mating differential (see below). Additionally, we emphasize that the measure capturing the underlying information of assortative mating is not the correlation itself, but rather a function of the square of the correlation coefficient, which takes values within the non-negative real number set [0, +∞).

### Fitness and sexual selection

Fitness, defined as an organism’s ability or propensity to survive and reproduce relative to others in the same population drives natural selection by leading to changes in gene frequencies (Hartl, 2020; Hedrick, 2005). Identifying natural selection relies on recognizing the patterns of change it produces in frequencies.

Consider a population with various types, each representing a set of individuals with specific properties (traits, genotypes, etc.). Let *N*_i_ be the number of individuals of type *i*, and suppose a selection episode changes this number to *N*’_i_. The absolute fitness, or growth rate, of type *i* during this episode is denoted as *W*_i_ = *N*’_i_/*N*_i_. To compare different types’ fitness fairly, we compute the relative fitness, *w*_i_, as the ratio of their absolute fitness to the mean population fitness. Thus, for a given type *i* with frequency *p*_i_, the new frequency *p*’_i_ after a selection episode will be given by (Bürger, 2000; Hansen, 2018)

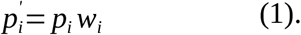

Recently, a simple theoretical framework, similar to the model presented in (1), has been developed to describe the influence of mating competition and mate choice on the emergence of sexual selection and assortative mating patterns (Carvajal-Rodríguez, 2018a, 2019, 2020). This theoretical framework has enabled the development of new statistical tests for detecting sexual selection and assortative mating. Moreover, by linking variation in mating ability (i.e., gamete access for fertilization) to the resulting frequency patterns, this framework facilitates the construction of predictive models and supports multi-model inference based on observed data patterns (Carvajal-Rodríguez, 2020).

In the following sections, I will briefly review and elucidate this theoretical framework, demonstrating its compatibility with Shuker and Kvarnemo’s (SK hereafter) definition of sexual selection (“any selection that arises from fitness differences associated with nonrandom success in the competition for access to gametes for fertilization”; Shuker and Kvarnemo 2021).

### Fitness for mating

We will focus on the change in frequencies due to fitness differences associated with access to gametes for fertilization. First, we must specify the concept of individual fitness for mating. Secondly, we will express for the discrete case, the SK definition of sexual selection in terms of information and then extend the definition to the continuous case.

Let *X* be a female character with a total of *k*_1_ discrete possible trait values and a male character *Y* with *k*_2_ discrete values. Characters *X* and *Y* can be the same or not. We define *M*_ij_ as the mutual ability or propensity to access gametes for fertilization between a female with value *i* for *X* and a male with value *j* for *Y*. Let *p*_1i_ be the population frequency of type *i* females and *p*_2j_ the frequency of type *j* males so that *q*_ij_ = *p*_1i_*p*_2j_ is the probability of random pairing of gametes from type *i* females with gametes from type *j* males for fertilization. The mean propensity is

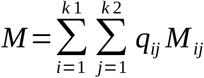

and the relative mutual propensity 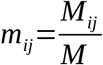.

The probability *q*’_ij_ of mating or gamete access for fertilization between any pair having such a combination of traits is (Carvajal-Rodríguez, 2018a)

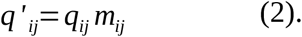

Equation (2) represents the expected frequencies of mating between types *i* and *j* as a function of the expected frequencies per random gamete encounter and the relative values of mutual propensity. The equation clearly resembles equation (1) for the change in frequencies because of selection. In fact, the absolute value *M*_ij_ can be expressed in a similar way to absolute fitness i.e., as the ratio of the number *N*’_ij_ of matings between types *i* and *j* after mating selection over the expected number *N*_ij_ if there were random mating.

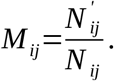

Our focus is on identifying the individual propensity to access gametes for fertilization, rather than the mutual propensity of pairs. We refer to this individual propensity as the fitness for mating. In this context, we define the absolute marginal propensity *M*_Fi_ of a female of type *i* as

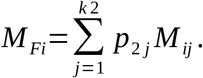

The relative marginal propensity or mating fitness *m*_Fi_ is obtained by dividing by the mean propensity *M*

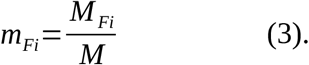

Similarly, the absolute marginal propensity *M*_Mj_ of a male of type *j* is

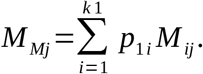

and the relative marginal propensity or mating fitness of males of type *j* is

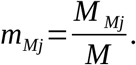

Thus, the relative mating fitness of a type is the average relative ability of individuals of that type to access gametes for fertilization.

While different mutual propensities shape mate choice, different mating fitness within a sex will produce a pattern of sexual selection (see Figure1 in Carvajal-Rodríguez, 2020). Sexual selection arises due to the unequal ability to access the gametes of the opposite sex for fertilization. Consequently, this sexual selection aligns perfectly with SK’s definition. Importantly, the concepts of mutual propensity and mating fitness, as defined, do not presuppose any particular mating preference function.

Having formalized the concepts of mutual propensity and mating fitness it is then possible to model mate competition and choice and the patterns of sexual selection and assortative mating they produce (see Carvajal-Rodríguez, 2018a, 2020).

The model in (2) facilitates the development of new statistical tests, based on information theory, to detect patterns of sexual selection and assortative mating (Carvajal-Rodríguez, 2018a). Moreover, it makes it possible to formalize the necessary and sufficient conditions for random mating in terms of mating propensities. Once these conditions are defined, we can generate different mating fitness scenarios that produce the intended patterns of sexual selection and assortative mating, allowing the application of powerful model inference techniques (Carvajal-Rodríguez, 2020).

The formalization of the mate choice and competition processes from the model presented in (2) have been carried out only for discrete traits and here we extend the mathematical framework to the continuous case.

### Sexual selection and assortative mating in quantitative traits

Consider two continuous traits in a population, *X* in females and *Y* in males, and let the mating probability distribution *Q*’(*x*,*y*) with probability density *q*’(*x*,*y*) represent the mating probability for pairs with values in the infinitesimal interval ([*x*,*x*+*dx*],[*y*,*y*+*dy*]). Alternatively consider the probability distribution *Q*(*x*,*y*) of random mating from the product of densities *q*(*x*,*y*)=*f*(*x*)*g*(*y*) of *X* and *Y*, respectively, where *f*(*x*) and *g*(*y*) are the population probability densities for traits *X* and *Y*, respectively.

Note that (2) still holds so that *q*’(*x*,*y*) depends on the density *q*(*x*,*y*) and the infinitesimal mutual mating propensity *m*_xy_ which is the relative mutual propensity to access gametes for fertilization between a female with value in the infinitesimal interval [*x*,*x*+*dx*] and a male with value in the infinitesimal interval [*y*,*y*+*dy*], so *q ‘* (*x*, *y*)=*m*_*xy*_ *q* (*x*, *y*).

In earlier work, Carvajal-Rodríguez (2018a) showed for discrete traits that non-random mating can be characterized as the Jeffreys divergence between the mating distribution, *Q*’, derived from mutual mating propensities, and the distribution of random mating, *Q*. The Jeffreys divergence is the information gain when changing from one distribution to another and vice versa. It is the sum of the two Kullback-Leibler divergences between *Q* and *Q*’, i.e., *J* = *KL*(*Q*’||*Q*)+*KL*(*Q*||*Q*’). For quantitative traits, it becomes

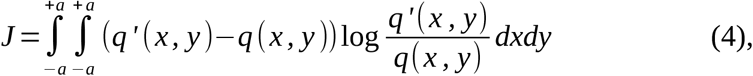

where *q*(*x*,*y*)=*f*(*x*)*g*(*y*). For simplicity and without loss of generality we assume that the range is the same for *X* and *Y* so that, *X*, *Y* ∈[−*a, a*].

If the information *J* is not significantly different from 0 we can assume that the distribution of matings *Q*’ is the same as that expected by random mating *Q* (Kullback, 1997). Note that the choice of *J* is not ad hoc; *J* is a measure of the information gain that arises when we measure the average change in the logarithm of mutual mating fitness (see Appendix A and Carvajal-Rodríguez, 2018a; Frank, 2012).

We will see that, just as in the discrete case (Carvajal-Rodríguez, 2018a), *J* can be decomposed into information due to sexual selection patterns in females and/or males (*J*_*PSS*_=*J*_*S1*_+*J*_*S2*_), assortative mating (*J*_*PSI*_) and an interaction term *E*. So, *J*=*J*_*PSS*_+*J*_*PSI*_+*E* (see below).

The distribution function of *X* in the population

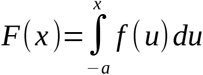

and *Y*

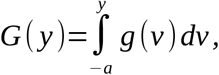

where *f*(*u*) and *g*(*v*) are the respective probability densities.

From the probability density *q*’(*x*,*y*), the marginal densities for *X* and *Y* in the matings are

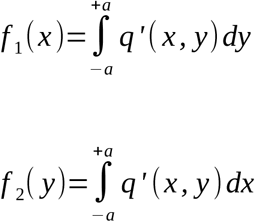

### Sexual selection

Sexual selection in females is measured as the difference between the phenotypic frequencies of females in matings and those in the population. In the continuous scenario, this can be expressed as the Jeffreys divergence between the population distribution of females and the marginal distribution of female matings.

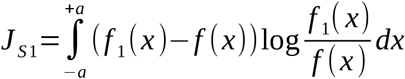

Similarly for males

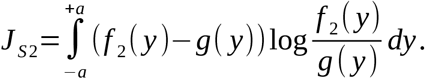

Recall that the pattern of sexual selection can be quantified by comparing the expected mating types resulting from random mating among mated individuals with those calculated using the population frequencies (Hartl and Clark, 1997; Rolán-Alvarez and Caballero, 2000). In the continuous scenario, this is formalized by the Jeffreys divergence

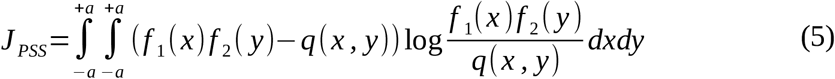

and because

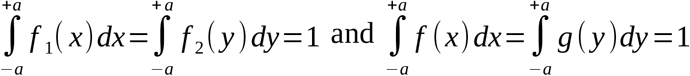

we have

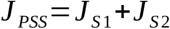

Therefore, similar to the approach applied in the discrete case (Carvajal-Rodríguez, 2018a), *J*_PSS_ quantifies sexual selection for quantitative traits as the change in phenotype distribution resulting from differential mating. The information captured in *J*_PSS_ is the sum of the sexual selection within each sex.

#### Normally distributed traits

Assume that *q*’(*x*,*y*) is a bivariate normal density and traits *X* and *Y* are normally distributed. Then, the marginal density function for the trait in females within matings is *f*_1_(x) ∼ *N*(µ_1_,σ_1_^2^), and in the population is *f*(*x*) ∼ *N*(µ_x_,σ_x_^2^). Similarly, the marginal density function for the trait in males within matings is *f*_2_(*y*) ∼ *N*(µ_2_,σ_2_^2^) and in the population is *g*(*y*) ∼ *N*(µ_y_,σ_y_^2^). Then, *J*_S1_ and *J*_S2_ can be expressed in closed form as (Belov and Armstrong, 2011; Kullback, 1997)

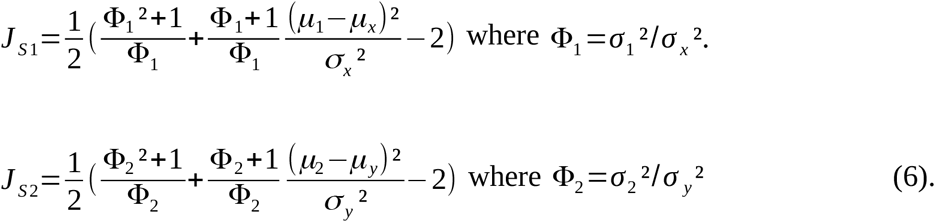

### Information statistics for sexual selection

With some sligth abuse of notation, consider now that *J*_*S*1_ and *J*_*S*2_ correspond to the values obtained using maximum likelihood estimators for the multivariate normal distribution. Then, for a random sample of *n* matings, *nJ*_S1_ (*nJ*_S2_) is asymptotically χ^2^ distributed with 2 degrees of freedom (Evren and Tuna, 2012; Kullback, 1997) under the null hypothesis that the females (males) within matings are distributed with mean µ_x_ (µ_y_) and variance σ_x_^2^ (σ_y_^2^) i.e. absence of sexual selection.

Note that in the above expressions, the term for the difference of the means is the square of the standardized sexual selection intensity (SSI) index (Caballero, 2020; Erlandsson and Rolán-Alvarez, 1998; Ng et al., 2019) so that

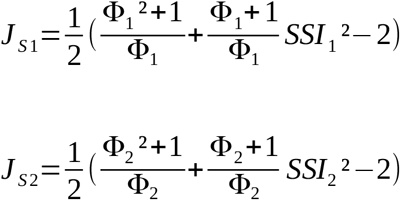

where *SSI*_1_ is the SSI index for females and *SSI*_2_ for males.

If we want to test only the mean value of the female trait *X* in the mating sample, we set Φ_1_=1 so that

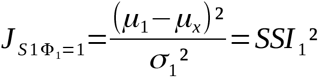

and with a sample size of *n* matings we have (*nJ*_S1Φ=1_)^0.5^ ∼ *t*_*n*-1_. Similarly, for the males we have (*nJ*_S2Φ=1_)^0.5^ ∼ *t*_*n*-1_ with

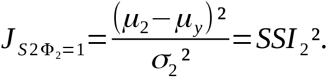

For large sample sizes both types of tests (χ^2^ or *t*) work similarly, for a smaller sample size the χ^2^ test might be slightly liberal and the *t*-test should be preferred.

More generally, the χ^2^ test should also detect differences in the variances, i.e., stabilizing or diversifying selection, or combinations of these forms of selection with directional selection (differences in means).

### Relationship of sexual selection information to other measures

There are several measures for the strength of sexual selection in quantitative traits (reviewed in Henshaw et al., 2016). Many of them are defined in terms of variation in reproductive success which is actually a much broader concept, representing an organism’s direct fitness (Shuker and Kvarnemo, 2021). Here we will make the connection in terms of mating success as a proxy for the mating fitnesses *m*_*F*_ and *m*_*M*_ as defined above. We will use male sexual selection as an example, but the approach for female sexual selection is analogous. Recall that we have defined the information due to the difference in the distribution of males within the matings with respect to the population distribution of males as

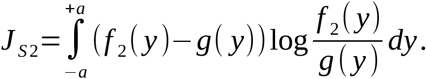

Also, the marginal density for *Y* in the matings was

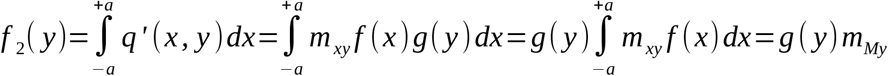

where *m*_My_ is the mating fitness of males with value for trait *Y* in the interval [*y*,*y*+*dy*].

The mean change in male trait caused by sexual selection is (Frank, 2012; Price, 1970; Taylor, 1996),

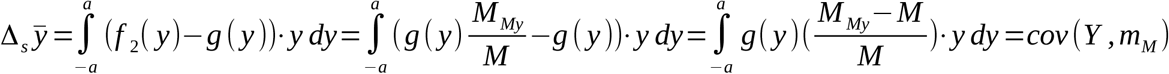

and if we take as character *Y* the logarithm of mating fitness, *L*_*M*_=log(*m*_*M*_), we get *J*_*S2*_, i.e.

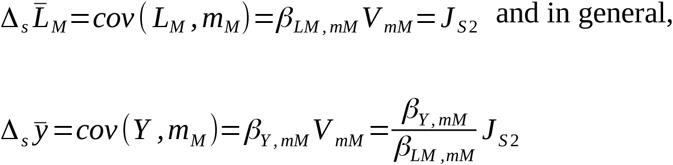

where *V*_*mM*_ is the variance in mating fitness and β_Y,mM_, β_*LM*,mM_ the regression coefficients of trait *Y* and log(*m*_*M*_) on fitness *m*_M_, respectively.

If changes are small and continuous (Frank, 2017, 2015) then β_LM,mM_ => 1 and

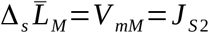

#### Opportunity for sexual selection, I_s_

For a given sex, the opportunity for sexual selection, *I*_*s*_, is equal to the variance in relative mating success (Wade and Arnold, 1980). Equating relative mating success with individual mating fitness, *m*_*M*_, for males, we get

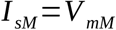

so for small continuous changes we have

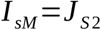

and in the general case

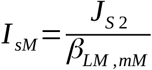

#### Standardized mating differential, s_m_

The mating differential (noted as *m*’ in Jones, 2009) is the covariance between standardized trait values and relative mating success, i.e.

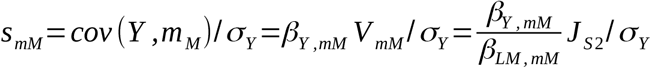

#### Bateman gradient and Jones index

The Bateman gradient β_ss_ is the slope of least-squares regression of relative reproductive success (*R*) on relative mating success (Arnold and Duvall, 1994; Bateman, 1948), i.e. for males β_ssM_ =β_R,mM_. The product of the Bateman gradient and the standardized mating differential is the standardized selection differential *s*’ (Jones, 2009) here *s’*_*M*_ for males, i.e. *s*’_*M*_=β_R,mM_ *s*_mM_. The Jones index is the maximum standardized sexual selection differential i.e., for males

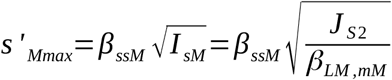

and similarly for females.

It is clear that the information measures, *J*_*S1*_ and *J*_*S2*_, developed here only refer to mating success, while the connection to reproductive success is part of other work that is ongoing.

### Assortative mating

A mating pattern can be described as the a posteriori deviation from random mating among mated individuals, resulting in positive assortative mating when similar phenotypes mate more frequently than expected, and negative assortative mating when the opposite occurs (Lewontin et al., 1968; Merrell, 1950; Spieth and Ringo, 1983).

Therefore, the assortative mating pattern corresponds to the divergence that compares the observed matings (*q*’) with those expected from random mating among mated individuals (*f*_1_(*x*)×*f*_2_(*y*))

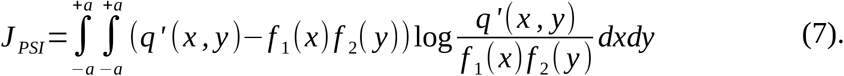

#### Normally distributed traits

As before, suppose that *q*’(*x*,*y*) is a bivariate normal density and that traits *X* and *Y* are normally distributed. The divergence for assortative mating in (7) is the Jeffreys divergence between the joint distribution and the product of the marginals and serves as a measure of the relationship between *X* and *Y* that can be expressed as a function of the correlation coefficient (see Appendix B and Kullback, 1997) so,

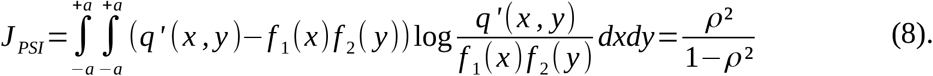

Thus, as long as the quantitative traits we are measuring are normally distributed and jointly normally distributed within mating pairs, the information measure of assortative mating is a function of the square of correlation coefficient that takes values within the set of nonnegative real numbers [0, +∞).

The statistical significance of *J*_PSI_>0 can be tested since *nJ*_PSI_ is asymptotically distributed as χ^2^ with 1 degree of freedom (Kullback 1997). To distinguish between positive and negative assortative mating, we only need to consider the sign of the Pearson correlation coefficient ρ.

### Total information

The difference *q*’(*x*,*y*)-*q*(*x*,*y*) in (4) can be partitioned as follows

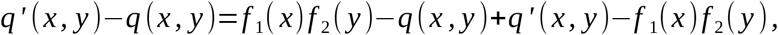

similarly, the logarithm

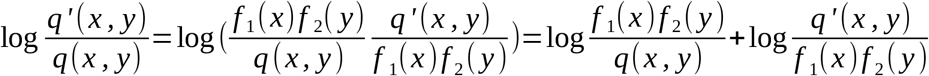

So, finally, the total divergence in (4) can be expressed as the sum of sexual selection (5) and assortative mating (7) divergences plus a factor *E*

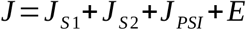

where (see Appendix A)

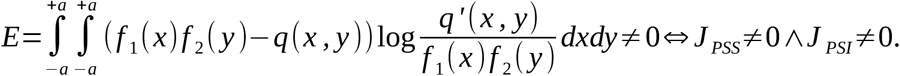

Carvajal-Rodríguez (2020) showed that from (2) we can define models where the mutual propensity to mate, *m*_*i*j_, is not multiplicative, that is, it is not equal to the product of the female and male marginals, *m*_*ij*_≠*m*_*Fi*_×*m*_*Mj*_, and these types of models correspond to mate choice models that generate specific mating patterns. Moreover, in addition to the assortative mating pattern, these models can also generate a frequency-dependent pattern of sexual selection. When this occurs, the increase in information reflected in both sexual selection and assortative mating patterns also includes a non-additive component that is captured by *E*. So, the *E* component only appears when both sexual selection (*J*_PSS_ ≠0) and assortative mating (*J*_PSI_ ≠0) patterns are present. It is usually a very low value.

For normally distributed traits, the total divergence *J* in (4) can be obtained in closed form and the non-additive component can be estimated by the difference of the additive components with the total (Appendix C)

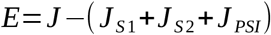

Furthermore, for normally distributed traits, it can be shown that the sum *J*_*S*1_+*J*_*S*2_ corresponds to equation (4) when the covariance in the pairings is 0 and *J*_*PSI*_ corresponds to equation (4) when the means and variances of both distributions (*Q*’ and *Q*) are equal (see Appendix C).

## Simulations

### Detection of sexual selection

We will utilize the program MateSim (Carvajal-Rodríguez, 2018b) to simulate the mating distribution for quantitative traits using a logistic model (Xie et al., 2015). This model represents an open-ended preference function, capable of generating a pattern of sexual selection in both females and males.

The simulation procedure consists of two main steps. First a population of size *N* is generated, comprising *N*/2 females and *N*/2 males. The individuals are randomly sampled from a standard normal distribution with a mean of 0 and a standard deviation of 1, *Z*(0,1). Second, a sample of *n* (*n*<*N*/2) matings is obtained as the individuals interact to form pairs for mating. The mating probability of a male *j* with phenotype *Z*_j_ of mating with any female *F* is evaluated as follows:

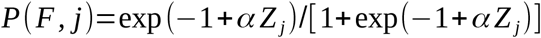

and the same applies for a female *i* with phenotype *Z*_i_ to mate with any male *M*.

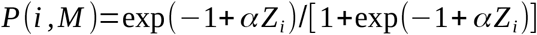

Therefore, the mating probability after encounter for a pair *i* × *j* will be

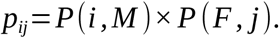

The parameter α (≥0) accounts for the strength of sexual selection, if α=0 there is random mating, the higher the value of α, the greater the strength of sexual selection.

The tests in (6) are able of distinguishing between random mating and sexual selection for different sample sizes and sexual selection intensities. Not surprisingly, the greater the sexual selection and the size of the sample, the greater the power of detection and the value of the information recovered by the *J*_S1_ and *J*_S2_ indexes (Table 1). For a large sample size, *n*=500, the correlation between the strength of sexual selection α and the *J* indexes was 0.99. For smaller sample size, *n*=50, the correlation was 0.97, indicating that *J*_S1_ and *J*_S2_ indices are good proxies of the strength of sexual selection.

**Table 1.** Mating under a logistic model. The parameter *α* accounts for the intensity of sexual selection. A value of *α* = 0 corresponds to random mating. Mating is with replacement. *N* is the population size and *n* the mating sample size. The quantitative trait distribution is *Z*(0,1). Number of runs: 10,000. % sig: The % of tests with *p*-value ≤0.05. The values in parentheses correspond to the average value of the statistic for the total number of significant runs.

| $N$ | $n$ | $\alpha$ | % sig. ($J_{s1}$) | %sig. ($J_{s2}$) |
| --- | --- | --- | --- | --- |
| $10^4$ | 500 | 0 | 5.3 (0.016) | 4.3 (0.016) |
| $10^4$ | 500 | 0.1 | 28.5.0 (0.019) | 30.0 (0.019) |
| $10^4$ | 500 | 0.25 | 96.6 (0.039) | 96.1 (0.038) |
| $10^4$ | 500 | 0.5 | 100 (0.126) | 100 (0.128) |
| $10^4$ | 500 | 1 | 100 (0.40) | 100 (0.41) |
| $10^4$ | 500 | 1.5 | 100 (0.710) | 100 (0.701) |
| $10^4$ | 500 | 2 | 100 (0.981) | 100 (0.985) |
| 500 | 50 | 0 | 5.0 (0.146) | 5.1 (0.141) |
| 500 | 50 | 0.1 | 9.5 (0.155) | 8.6 (0.15) |
| 500 | 50 | 0.25 | 25.9 (0.168) | 28.2 (0.179) |
| 500 | 50 | 0.5 | 64.3 (0.215) | 66.3 (0.219) |
| 500 | 50 | 1 | 99.9 (0.586) | 99.5 (0.476) |
| 500 | 50 | 1.5 | 100 (0.845) | 100 (0.918) |
| 500 | 50 | 2 | 100 (1.107) | 100 (1.044) |

### Detection of assortative mating: *J*_PSI_ versus correlation

We will utilize the program MateSim (Carvajal-Rodríguez, 2018b) to simulate the mating distribution for quantitative traits using a FND Gaussian function (Carvajal-Rodríguez and Rolán-Alvarez, 2014). The FND function with parameter *C*>0 generates positive assortative mating patterns that can be measured by the Pearson correlation coefficient ρ and also by *J*_PSI_ as given in (8).

The simulation procedure is as before but the preference is evaluated following the FND function

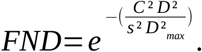

If *C*=0 the mating is random, if *C*>0 there is mate choice that generates positive assortative mating. The parameter *D* represents the phenotypic difference between male and female in the pair. If the *D* is zero the preference is the maximum. The parameter *s* is the tolerance to mate with a partner that does not correspond to the preference optimum. i.e. *s*= 0.08 implies eight times more tolerance for non-optimal mating than *s*=0.01. *D*_max_ is the maximum phenotypic difference in the population between any two individuals, regardless of sex. For a phenotypic difference *D*/*D*_max_ between a pair, the strength of the choice depends on the ratio between the choice and tolerance parameters, *C*/*s*.

For a sample of *n* matings, the significance of ρ can be computed as 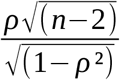 that follows a *t*-distribution with *n*-2 degrees of freedom. We have already seen that *nJ*_*PSI*_ is asymptotically distributed as χ^2^ with 1 degree of freedom.

In Table 2 both types of tests are performed for mating distributions generated under different choice and tolerance values. When the population and sample size are large, 10,000 and 500, respectively, from choice values of 0.2, both the correlation and the *J*_PSI_ have greater than 97% power to detect assortative mating. The difference is that the correlation value already reaches its maximum for choice values of 0.4.

**Table 2.** One sex is choosy. Choice implies preference for similarity. Mating is with replacement. *N* is the population size and *n* the sample size of matings. The quantitative trait distribution is *Z*(0,1). Number of runs: 10,000. % sig: The % of tests with *p*-value ≤0.05. The values in parentheses correspond to the average value of the statistic for the total number of significant runs.

| $N$ | $n$ | *Choice* | $s$ | % sig. ( $\rho$ ) | %sig. ($\rho^2/(1-\rho^2)$ ) |
| --- | --- | --- | --- | --- | --- |
| $10^4$ | 500 | 0 | -- | 5.1 (0.01) | 5.2 (0.01) |
| $10^4$ | 500 | 0.2 | 0.08 | 97.9 (0.18) | 97.9 (0.03) |
| $10^4$ | 500 | 0.4 | 0.08 | 100 (0.55) | 100 (0.43) |
| $10^4$ | 500 | 0.8 | 0.08 | 100 (0.85) | 100 (2.60) |
| $10^4$ | 500 | 1 | 0.08 | 100 (0.89) | 100 (3.72) |
| $10^4$ | 500 | 0.2 | 0.01 | 100 (0.95) | 100 (9.3) |
| $10^4$ | 500 | 0.4 | 0.01 | 100 (0.99) | 100 (59.0) |
| $10^4$ | 500 | 0.8 | 0.01 | 100 (1.0) | 100 (194.0) |
| $10^4$ | 500 | 1 | 0.01 | 100 (1.0) | 100 (413.4) |
| 500 | 50 | 0 | -- | 5.1 (0.03) | 6.2 (0.12) |
| 500 | 50 | 0.2 | 0.08 | 38.6 (0.37) | 41.7 (0.16) |
| 500 | 50 | 0.4 | 0.08 | 99.95 (0.61) | 99.97 (0.64) |
| 500 | 50 | 0.8 | 0.08 | 100 (0.90) | 100 (4.59) |
| 500 | 50 | 1 | 0.08 | 100 (0.93) | 100 (6.68) |
| 500 | 50 | 0.2 | 0.01 | 100 (0.96) | 100 (13.54) |
| 500 | 50 | 0.4 | 0.01 | 100 (0.99) | 100 (56.68) |
| 500 | 50 | 0.8 | 0.01 | 100 (1.0) | 100 (183.08) |
| 500 | 50 | 1 | 0.01 | 100 (1.0) | 100 (403.27) |

When the population and sample sizes are smaller, specifically 500 and 50, respectively, we observe a few noteworthy points. Firstly, it is not surprising that *J*_PSI_ tends to be somewhat liberal in these cases, as the test *nJ*_PSI_ is only asymptotically distributed as χ^2^, and its behavior improves with higher sample sizes. Secondly, both methods demonstrate approximately 40% power when the choice is 0.2 and the tolerance is 0.08.

However, they exhibit almost 100% power in other cases, such as when the tolerance is 0.01 or the choice is from 0.4 and beyond. The difference between ρ and *J*_PSI_ is again the saturation of the value of ρ as the strength of choice increases.

*J*_*PSI*_ achieves higher resolution when the strength of choice is sufficiently high to yield correlation values close to 1. For instance, as demonstrated in Table 2, a choice *C* of 0.2 with a low tolerance of *s*=0.01 produces a mating correlation of 0.95. Consequently, for values exceeding *C*>0.2 under a tolerance of 0.01, *J*_*PSI*_ will more effectively discriminate between varying degrees of assortative mating.The greater the sample size and the choice strength, the greater the power to detect assortative mating. In Figure 1, the correlation of the values in Table 1 i.e., between the choice strength (*C*/*s*) and both statistics, can be appreciated. For the ρ statistic the correlation with choice strength is 0.6 for both large and small sample size, for *J*_PSI_ the correlation with the choice strength is 0.97.

**Figure 1.**
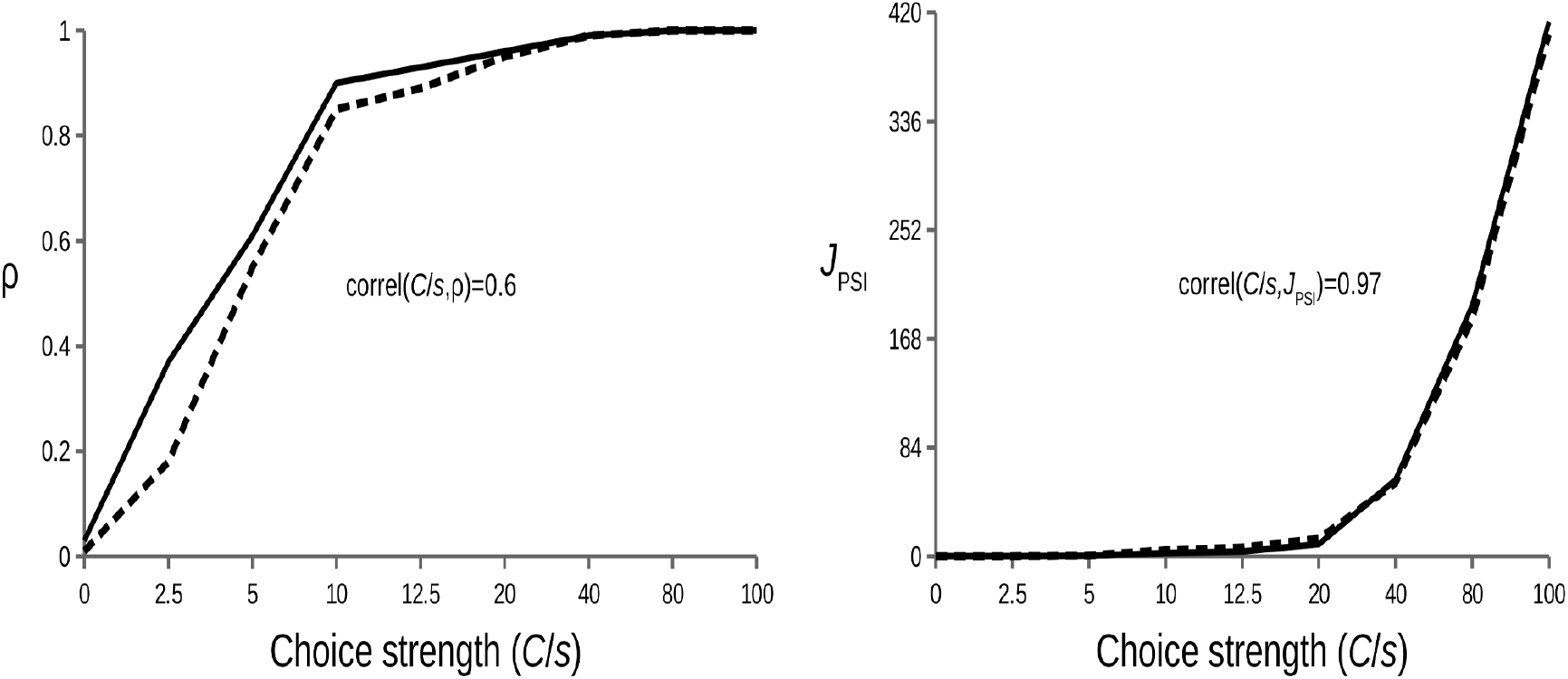
Graphic representation of values of the rho statistics (left panel) and *J*_*PSI*_ (right panel) with respect to the strength of the choice (*C/s*). The value of the correlation between each statistic and the strength of the choice is given. Continuous line: population size *N*=10,000, sample size *n*=500. Dashed line: *N*=500, *n*=50. The values of ρ and *J*_*PSI*_ are those in Table 1 ordered by increasing *C*/*s*.

The results were almost identical (Appendix D Table S1) when we consider a bias for the mating preference, i.e. the choosy males preferring females 1.5 times bigger than themselves (Lau et al., 2021).

We were also interested in evaluating the performance of the *J*_PSI_ index across different functions and mating scenarios in which there was competition for mating (which generates sexual selection patterns) but no mate choice (no assortative mating pattern). To achieve this, we employed the logistic function with α =2 which implies sexual selection without mate choice. The mating process was implemented with replacement, where each individual can mate more than once, that is, the availability of individuals is not affected by matings that have already occurred. Additionally, mating without replacement was also considered, where the proportion of individuals available for mating must be updated after each mating event. In the case of no replacement, the encounters can be individual or massive (Gimelfarb, 1988). Individual encounters imply that only one encounter occurs at a time, while when mating occurs with mass encounters, more than one pair can be formed simultaneously (see Appendix D Table S2). In the case of large population and sample sizes, both ρ and *J*_PSI_ exhibited the expected false positive rate of 5%. However, with lower sample sizes, *J*_PSI_ showed a slightly liberal false positive rate of 6%.

We also simulate a mate choice scenario for a bimodal population where two more or less extreme phenotypes coexist (Appendix D Table S3). The power for the detection of assortative mating was similar to the unimodal case and the correlation of the statistics with the strength of choice was greater the larger the sample size, being about 0.58-0.65 for ρ and 0.95-0.97 for *J*_PSI_ (Figure 2).

**Figure 2.**
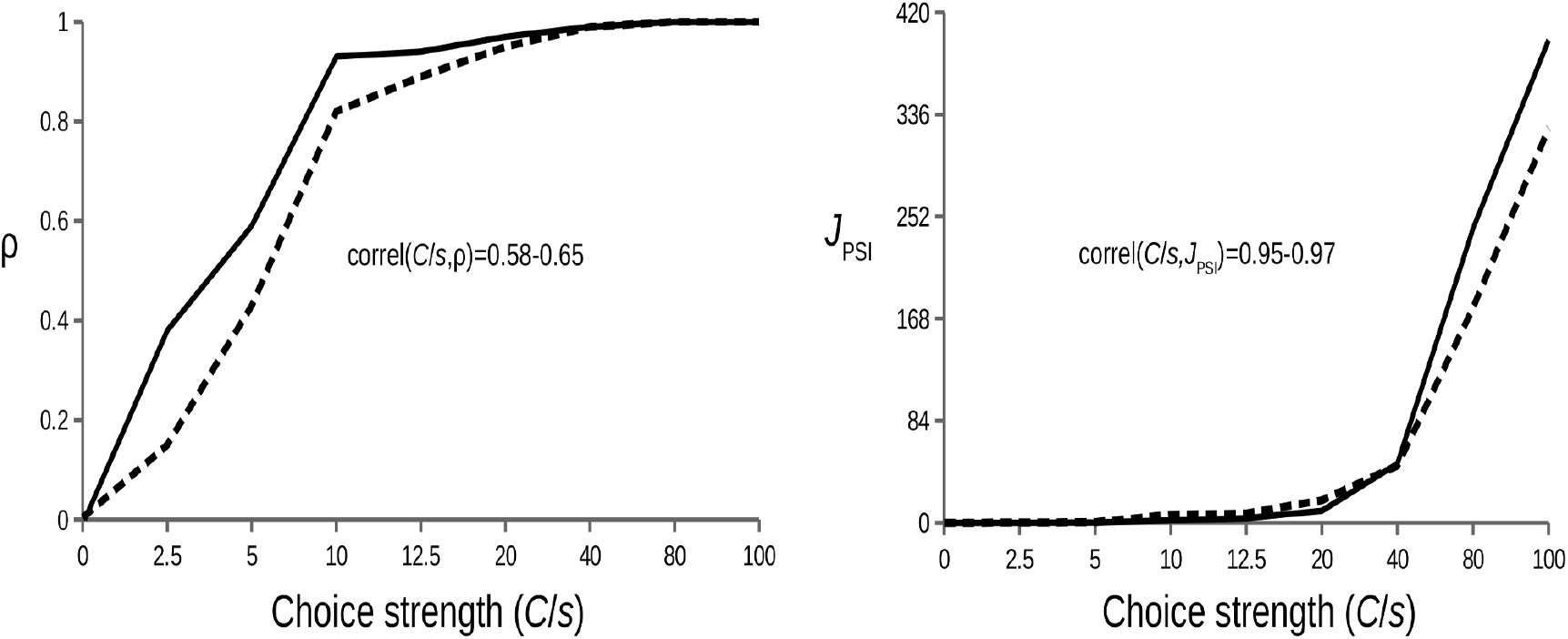
Graphic representation of values of the rho statistics (left panel) and *J*_*PSI*_ (right panel) with respect to the strength of the choice (*C*/*s*). Bimodal population with mixture proportion π=0.5, µ_2_-µ_1_=3 and σ_1_+σ_2_ = 2. Continuous line: population size *N*=10,000, sample size *n*=500. Dashed line: *N*=500, *n*=50. The values of ρ and *J*_*PSI*_ are those in Table S3 ordered by increasing *C*/*s*.

Also, we examined how being choosy in one or both sexes affects the ρ and *J*_PSI_ indexes and found no differences (compare Table S4 in Appendix D with Table 1).

Lastly, we check that *J*_*PSI*_ still performs well for smaller choice strength values (Table S5 and Figure S1 in Appendix D).

### Example of application

*Echinolittorina malaccana* is a marine intertidal snail commonly found on rocky shores throughout the Indo-West Pacific region (Reid, 2007). The primary mechanism of mate choice in this species appears to be male preference for females slightly larger than themselves. Assortative mating for size, measured by pairwise correlation, has been estimated to be around 0.5 (López-Cortegano et al., 2020; Ng et al., 2019). In a previous study, size (shell length, a continuous trait) has been transformed into a qualitative trait (discrete size classes) enabling the use of multimodel inference techniques. These analyses provided deeper insights into mate choice and confirmed that the most suitable model involves mate choice with bias, resulting in both assortative mating and sexual selection for size (Lau et al., 2021).

We will use the same onshore data from mating pairs of *E. malaccana* used in Lau et al. (Lau et al., 2021) and available from Ng et al. (Ng et al., 2019; Rolán-Alvarez et al., 2021) and Rolán-Alvarez (Rolán-Alvarez et al., 2021) to apply the new indices *J*_PSS_ and *J*_PSI_ directly to the shell length trait (Table 3).

**Table 3.** Analysis of *Echinolittorina malaccana* data.

| Test | N | n | $J_{S1}$ (pval) | $J_{S2}$ (pval) | $\rho$ (pval) | $J_{PSI}$ (pval) |
| --- | --- | --- | --- | --- | --- | --- |
| Chi<sup>2</sup> | 876 | 228 | 0.11 (<10<sup>-5</sup>) | 0.03 (0.031) | 0.57 (<10<sup>-5</sup>) | 0.49 (<10<sup>-5</sup>) |

Data from Ng et al 2019.

Sexual selection was observed in both sexes, although it appeared to be weaker in males. This finding aligns with previous research, indicating that sexual selection in *E. malaccana* females is likely driven by male preference for slightly larger females, while in males, it may result from male-male competition (Lau et al., 2021; Ng et al., 2019). As for assortative mating, both the Pearson correlation coefficient and the new *J*_PSI_ index supported and validated previous findings.

## Discussion

There is a vast amount of literature exploring mate choice, competition, assortative mating, and sexual selection as influential forces in the evolution of traits that may not always be directly favoured by natural selection (Andersson, 1994; Darwin, 1871; reviewed in Lindsay et al., 2019). These concepts are traditionally grouped under the term ‘theory of sexual selection’. Sexual selection theory stands as one of the most active areas of evolutionary research, engaging diverse fields and inference methodologies, given its relevance in population genetics, evolutionary ecology, animal behavior, sociology, and psychology. Two recent important reviews were published on the concept and definition of sexual selection (Alonzo and Servedio, 2019; Shuker and Kvarnemo, 2021). Alonzo and Servedio discuss the challenges in achieving a consensus definition, while Shuker and Kvarnemo propose an integrative definition for sexual selection.

In this study, we demonstrate that the SK definition of sexual selection can be effectively applied within a modeling framework that utilizes information theory to describe mate choice and competition processes, leading to patterns of sexual selection and assortative mating. Under the information theory formalism, these patterns of sexual selection and assortative mating are described in terms of Jeffreys divergence.

Importantly, this novel framework, initially developed for discrete traits, is general and doesn’t assume a priori whether the choice is exercised by males, females, or both, nor does it make assumptions about the sex that competes. Furthermore, it is agnostic concerning the genetic basis or evolutionary mechanisms underlying complex mate choice and competition processes. The informational framework starts with a concept of mutual mating fitness or mutual mating propensity (sensu Carvajal-Rodríguez, 2020, 2018a), where the marginal values correspond to individual mating fitness. These mating fitnesses allow us to measure the pattern of sexual selection according to the SK definition.

In addition, we have extended the informational framework to encompass quantitative traits. For both discrete and quantitative traits, sexual selection is detected by comparing the distribution of mating phenotypes with that of the overall population. Similarly, assortative mating is detected by comparing, within mating couples, the observed joint distribution of mating traits to that expected under random mating.

When the traits are normally distributed, the sexual selection indexes *J*_S1_ (females) and *J*_S2_ (males) can be expressed as a function of the standardized sexual selection intensity index and the ratio of sample and population variances (6). Simulations have shown that *J*_S_ indexes have a correlation of 0.99 with the strength of sexual selection.

When we compare sexual selection information indices with other measures of sexual selection for quantitative traits, we see that if we consider mating success as a proxy for mating fitness, the sexual selection information index coincides with the opportunity for sexual selection (Wade and Arnold, 1980), and from here we can establish the relationship with other measures such as the standardized mating differential (Jones, 2009). Therefore, in this work, we have demonstrated that the average change in a quantitative trait resulting from differential mating fitness corresponds to the increase in information quantified by the sexual selection information indexes *J*_*S*1_ and *J*_*S*2_.

Additionally, we have established a connection between *J*_*S*1_+*J*_*S*2,_ and the statistic for discrete traits *PSS* (Rolán-Alvarez and Caballero, 2000), as well as with measures of quantitative traits based on mating success.

Furthermore, the assortative mating index *J*_PSI_, when the traits are normally distributed, is a transformation of ρ and has as much statistical power as ρ, with the added advantage of being highly correlated with the strength of mate preference, which is not generally true for the correlation coefficient (Clancey et al., 2022; Fernández-Meirama et al., 2022).

Finally, we applied the quantitative trait information indices developed in this work to a previously published dataset on size assortative mating and sexual selection (Lau et al., 2021; Ng et al., 2019). We confirmed previous findings and validate the quantitative approach for the new indices of sexual selection and assortative mating.

### Limitations of the model

We have demonstrated that the concept of mating fitness derived from the *q*′=*qm* model aligns perfectly with S&K’s definition of sexual selection. This definition encompasses postcopulatory processes, which are included under the term ‘access to gametes for fertilization’ consistent with our concept of differential mating fitness. However, the definition and measurement of sexual selection are different concepts, and the same applies to the definition of mating fitness and its measurement. Thus, when mating frequencies are used, regardless of whether fertilization occurs or not, as a proxy to estimate mutual and individual mating fitnesses, we are only focusing on precopulatory mechanisms of sexual selection.

Similarly, when discussing the measurement of sexual selection, we have compared the indices *J*_*S1*_ and *J*_*S2*_ with other measures of sexual selection and considered mating success as a proxy for mating fitness to equate *J*_*S*_ with the opportunity for sexual selection for continuous and small changes. However, as we have already mentioned, if the proxy used for mating fitness is based on the ratio of observed to expected mating frequencies, these estimates may not be adequate for various scenarios, so the connection with variances in mating or reproductive success may not be accurate, or the variances may not be due to sexual selection (Henshaw et al., 2016). Therefore, regarding the presented model and its interpretation from information theory, the limitations are not so much of the model but rather the difficulty in estimating mating fitness from certain characteristics.

### Concluding remark

In conclusion, the application of information theory to describe patterns of sexual selection and assortative mating (Carvajal-Rodríguez, 2020, 2019, 2018a), has paved the way for extending these developments to quantitative traits. This study demonstrates that sexual selection and assortative mating can be effectively quantified as information gain relative to the random mating pattern. Just like the previous developments for discrete traits, the results obtained with quantitative traits indicate that employing information theory to describe mating processes and patterns holds great promise as a valuable approach to understanding both the causes and effects of sexual selection.

## Acknowledgements

Dedicated to my sister Paci. I thank to Emilio Rolán-Alvarez and two anonymous reviewers for their useful comments. This work was supported by Xunta de Galicia (Grupo de Referencia Competitiva, ED431C 2020/05), Ministerio de Ciencia e Innovación (PID2022-137935NB-I00) and by Fondos FEDER. Funding for open access charge: Universidade de Vigo/CISUG.

## Appendix

### A. General formula for non-random mating information

Consider two continuous traits, *X* in females and *Y* in males from a population and let the probability distribution *Q*’(*x*,*y*) with probability density *q*’(*x*,*y*) that represents the mating probability for pairs with values in the infinitesimal interval ([*x*,*x*+*dx*],[*y*,*y*+*dy*]). Consider alternatively the probability distribution *Q*(*x*,*y*) of random mating with the product of densities *q*(*x*,*y*)=*f*(*x*)*g*(*y*) from *X* and *Y*, respectively.

Let *x* and *y*, be phenotypes in a mating pair, and let *Z*_*xy*_ be any function of the traits (*x*,*y*). Now, choose the function *L*_*xy*_=log(*m*_*xy*_) so that *Z*_*xy*_=*L*_*xy*_. Then, the average change in *L* caused by non-random mating is given by Jeffreys divergence (Carvajal-Rodríguez, 2018a; Frank, 2012)

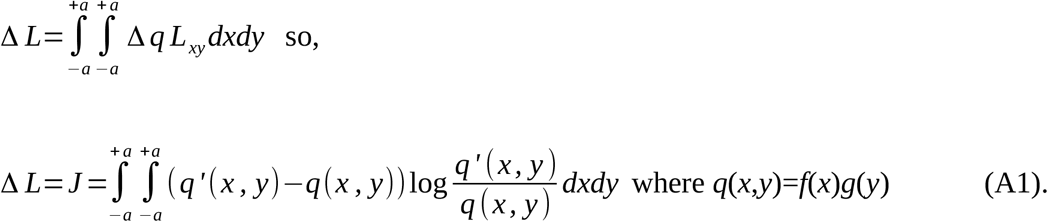

However, *Z* can be any function of the pairs *(x*,*y*). For example, *Z* may be a linear function of the display (*δ*) by the courter (sensu Rosenthal, 2017) and the preference (π) of the chooser e.g., Z_xy_=*aδ+bπbπ*, where *a* and *b* are ad hoc weight functions. If both traits *x* and *y* are size and only one sex chooses (e.g. females), then *δ=y, ay, a*=1, and *b*=0, so Z_xy_=*y*. However, if there is a preference for males of similar size, then it could be, *δ=y, a0, b*=1, *π=y, a*1-|x-y|/*D*_*max*_, where *D*_*max*_ is the maximum possible difference between *x* and *y*.

The relationship of J with the average change in Z, caused by mutual mating fitness is

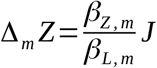

where β_Z,m_ is the regression of *Z* on the actual mutual mating propensity *m* and β_L,m_ is the regression of log(*m*) on *m*.

With respect to *J*, the difference *q*’(*x*,*y*)-*q*(*x*,*y*) can be partitioned as follows

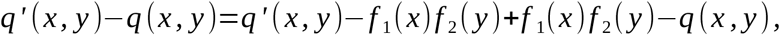

where *f*_1_(*x*) and *f*_2_(*y*) are the marginal densities for *X* and *Y*

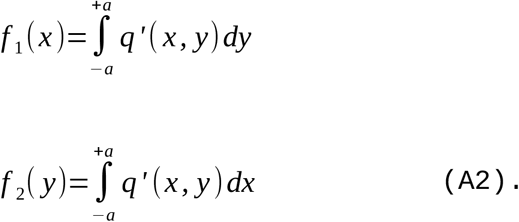

Similarly, the logarithm

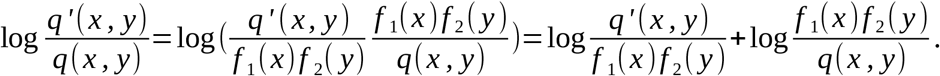

Now note that the densities *f*(*x*) and *g*(*y*) depend on the population distribution, while the marginal densities *f*_1_(*x*) and *f*_2_(*y*) depend on the mating distribution so they are not necessarily equal. Then we have

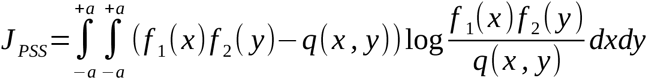

and

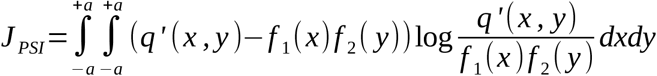

therefore, as in the discrete case, the measure of non-random mating can be divided into

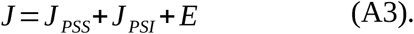

The component *E* is

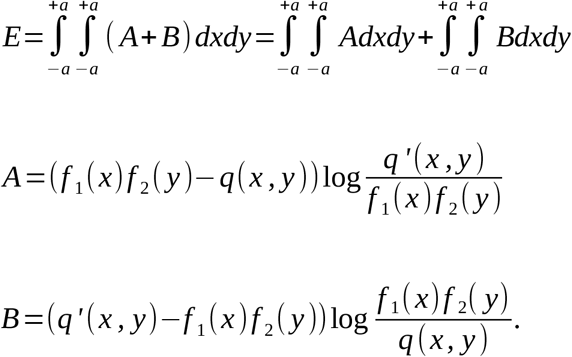

We will show that

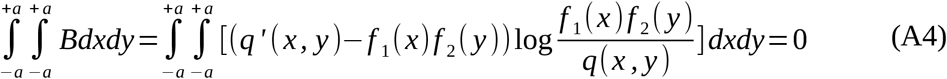

so that,

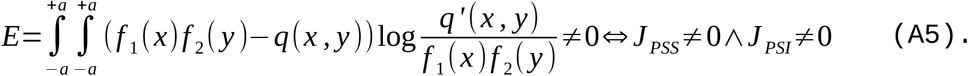

Now, first recall that *q*(*x*,*y*)=*f*(*x*)*g*(*y*) and then

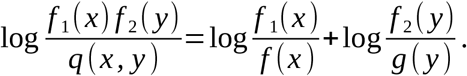

Also,

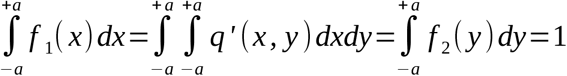

so that,

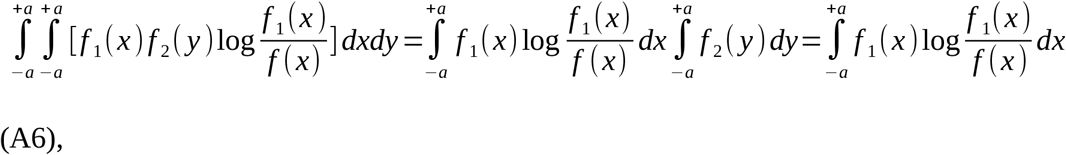

and similarly

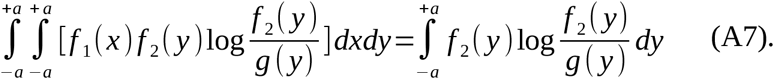

Also using the marginal density definitions from (A2)

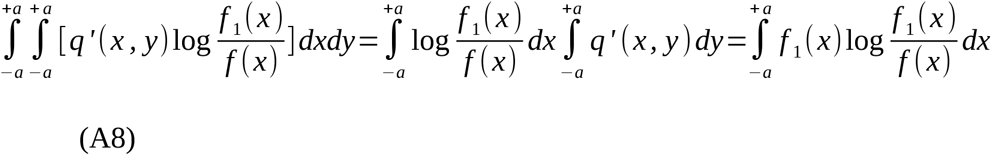

and

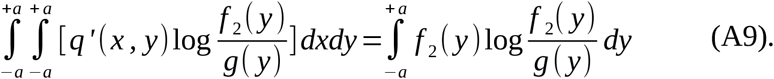

Finally, noting that (A6) equals (A8) and (A7) equals (A9) and rearranging and substituting in (A4) we obtain

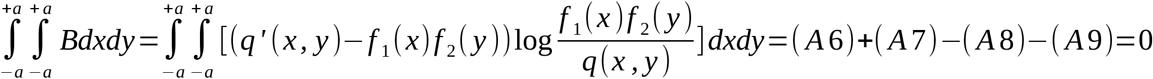

### B. Assortative mating with normally distributed traits

The general formulation is

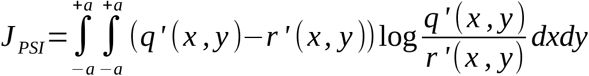

where *r*’(*x*,*y*)= *f*_1_(*x*)*f*_2_(*y*)

If *q*’ is bivariate normal and *f*_1_ and *f*_2_ are normally distributed we have

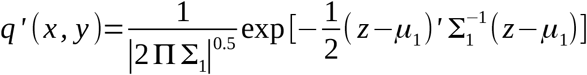

where Σ_1_ is the variance-covariance matrix (see below), |Σ_1_| is the determinant, *z* is the column vector of the two variables *x*,*y* and *z*’ is the row vector of *x*,*y* and *μ*_1_ is the column vector (*μ*_1x_,*μ*_1y_).

Similarly, the product of the two marginals is

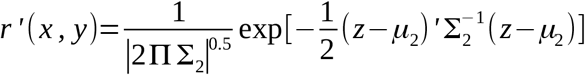

where *μ*_2_ is the column vector (*μ*_2x_,*μ*_2y_).

Note that in our case, using males and females from matings (joint distribution versus random mating distribution obtained from mating data) implies that mean and variances are equal between distributions (μ_1_=μ_2_, σ_1_=σ_2_), i.e. μ_1x_=μ_2x_, μ_1y_=μ_2y_ and σ_1x_=σ_2x_, σ_1y_=σ_2y_.

Let

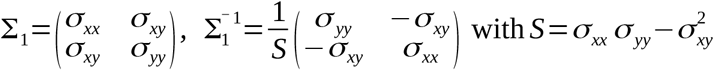

and

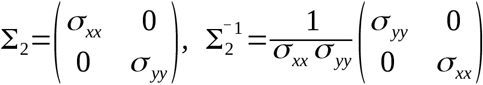

In this case the Kullback-Leibler divergences are (c.f. Kullback, 1997 (1.6) p. 190)

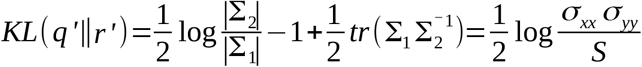

and

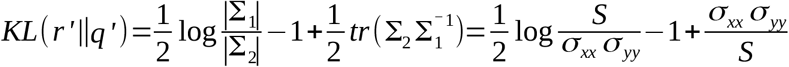

where *tr* is the trace (sum of diagonal elements) of a square matrix.

Therefore,

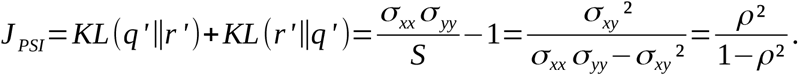

### C. Total *J* with normally distributed traits

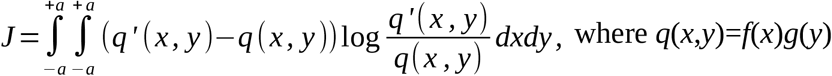

If *q*’ is *N*(μ_1_, Σ_1_) and *q N*(μ_2_, Σ_2_) where Σ_i_ is the variance-covariance matrix, we have that the *KL* divergence has the following closed form (Pardo, 2018)

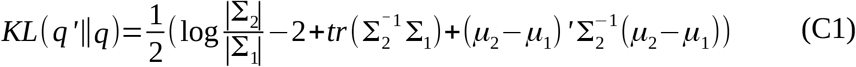

where *tr* is the trace (sum of diagonal elements) of a square matrix, |Σ| is the determinant, *μ*_*i*_ is the column vector (μ_ix_, μ_iy_), *μ*’ is the row vector.

Therefore,

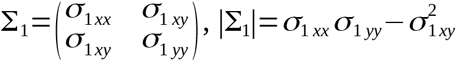

and

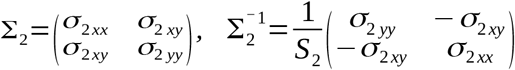

where

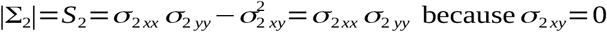

Now, let

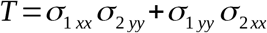

then

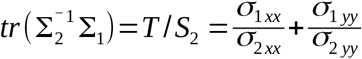

and,

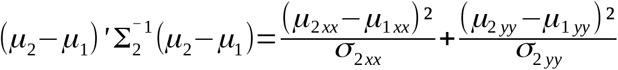

The other divergence is

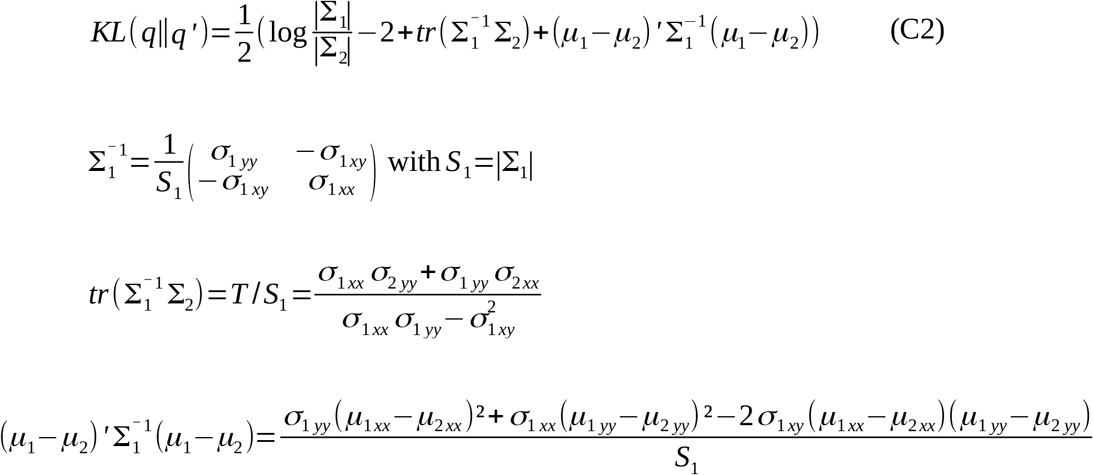

Finally,

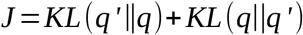

with *nJ* asymptotically χ^2^ distributed with 5 degrees of freedom where *n* is the sample size of matings.

#### C.1 Mating covariance greater than 0 with equal means and variances

Note that from (C1) and (C2), if we assume that the means and variances between the two distributions are the same, i.e. µ_1_=µ_2_, σ_1xx_=σ_2xx_, σ_1yy_=σ_2yy_ but σ_1xy_>0 and σ_2xy_=0 then

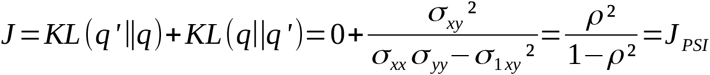

which is the assortative mating component without sexual selection (*J*_*PSI*_) obtained in the Appendix B.

#### C.2 No covariance within matings

On the other side, if we assume σ_1xy_=σ_2xy_=0 then from C1 and C2 we get

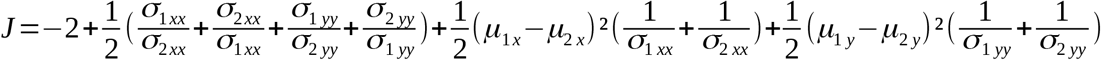

which can be expressed as

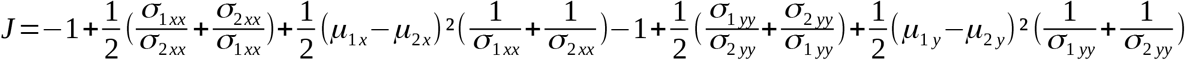

and by taking Φ_1_ =*σ*_1 *xx*_ / *σ*_2 *xx*_ and Φ_2_=*σ*_1 *yy*_ / *σ* _2 *yy*_ and noting that

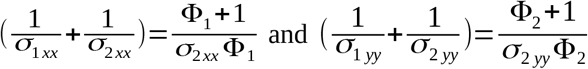

then
*J*=*J*_*S*1_+*J*_*S*2_ as defined in (6), which is sexual selection without assortative mating.

#### C.3 No covariance and equal means

Even if we assume only differences in variances, we obtain the information gain due to the pattern of sexual selection as

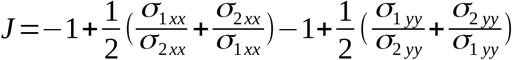

that corresponds to (6) when the means are assumed to be equal.

#### D. Simulation results tables

**Table S1.** One sex is choosy. Choice implies preference for similarity with a bias of 1.5. Mating is with replacement. *N* is the population size and *n* the sample size of matings. The quantitative trait distribution is *Z*(0,1). Number of runs: 10,000. % sig: The % of tests with *p*-value ≤0.05. The values in parentheses correspond to the average value of the statistic for the total number of significant runs.

| $N$ | $n$ | Choice | $s$ | % sig. ( $\rho$ ) | %sig. ($\rho^2/(1-\rho^2)$ ) |
| --- | --- | --- | --- | --- | --- |
| $10^4$ | 500 | 0 | -- | 4.8 (0.004) | 4.9 (0.01) |
| $10^4$ | 500 | 0.2 | 0.08 | 99.9 (0.22) | 99.9 (0.05) |
| $10^4$ | 500 | 0.4 | 0.08 | 100 (0.54) | 100 (0.41) |
| $10^4$ | 500 | 0.8 | 0.08 | 100 (0.87) | 100 (3.19) |
| $10^4$ | 500 | 1 | 0.08 | 100 (0.89) | 100 (3.99) |
| 500 | 50 | 0 | -- | 5 (-0.025) | 5.9 (0.12) |
| 500 | 50 | 0.2 | 0.08 | 44.7 (0.37) | 48.2 (0.17) |
| 500 | 50 | 0.4 | 0.08 | 100 (0.67) | 100 (0.90) |
| 500 | 50 | 0.8 | 0.08 | 100 (0.84) | 100 (2.54) |
| 500 | 50 | 1 | 0.08 | 100 (0.90) | 100 (4.43) |
| 500 | 50 | 0.2 | 0.01 | 100 (0.97) | 100 (64.72) |
| 500 | 50 | 0.4 | 0.01 | 100 (0.99) | 100 (41.97) |
| 500 | 50 | 0.8 | 0.01 | 100 (1) | 100 (115.87) |
| 500 | 50 | 1 | 0.01 | 100 (1) | 100 (3935.1) |

**Table S2.** Mating under a logistic model with α=2. There is no preference for similarity. *N* is the population size and *n* the sample size of matings. The quantitative trait distribution is *Z*(0,1). Number of runs: 10,000. % sig: The % of tests with *p*-value ≤0.05. The values in parentheses correspond to the average value of the statistic for the total number of significant runs.

| $N$ | $n$ | Mating | % sig. ( $\rho$ ) | %sig. ($\rho^2/(1-\rho^2)$ ) |
| --- | --- | --- | --- | --- |
| $10^4$ | 500 | replacement | 5.3 (0.003) | 5.4 (0.01) |
| 10<sup>4</sup> | 500 | individual | 4.9 (0.003) | 5.0 (0.01) |
| 10<sup>4</sup> | 500 | mass-encounter | 4.7 (0.004) | 4.8 (0.01) |
| 500 | 50 | replacement | 5.0 (-0.01) | 6.0 (0.12) |
| 500 | 50 | individual | 5.2 (0.02) | 6.3 (0.12) |
| 500 | 50 | mass-encounter | 5.4 (0.03) | 6.4 (0.12) |

**Table S3.** Bimodal populations with mixture proportion π=0.5, µ_2_-µ_1_-=3 and σ_1_+σ_2_ = 2. One sex is choosy. Choice implies preference for similarity. Mating is with replacement. *N* is the population size and *n* the sample size of matings. The quantitative trait distribution is *Z*(0,1). Number of runs: 10,000. % sig: The % of tests with *p*-value ≤0.05. The values in parentheses correspond to the average value of the statistic for the total number of significant runs.

| $N$ | $n$ | *Choice* | $s$ | % sig. ( $\rho$ ) | %sig. ($\rho^2/(1-\rho^2)$ ) |
| --- | --- | --- | --- | --- | --- |
| 10<sup>4</sup> | 500 | 0 | -- | 5.1 (0.004) | 5.2 (0.01) |
| 10<sup>4</sup> | 500 | 0.2 | 0.08 | 90 (0.15) | 90.1 (0.03) |
| 10<sup>4</sup> | 500 | 0.4 | 0.08 | 100 (0.43) | 100 (0.22) |
| 10<sup>4</sup> | 500 | 0.8 | 0.08 | 100 (0.82) | 100 (1.98) |
| 10<sup>4</sup> | 500 | 1 | 0.08 | 100 (0.89) | 100 (3.79) |
| 10<sup>4</sup> | 500 | 0.2 | 0.01 | 100 (0.95) | 100 (9.99) |
| 10<sup>4</sup> | 500 | 0.4 | 0.01 | 100 (0.99) | 100 (48.67) |
| 10<sup>4</sup> | 500 | 0.8 | 0.01 | 100 (1.0) | 100 (242.35) |
| 10<sup>4</sup> | 500 | 1 | 0.01 | 100 (1.0) | 100 (397.16) |
| 500 | 50 | 0 | -- | 5.1 (-0.01) | 6.1 (0.12) |
| 500 | 50 | 0.2 | 0.08 | 43.2 (0.38) | 46.2 (0.18) |
| 500 | 50 | 0.4 | 0.08 | 99.7 (0.59) | 99.7 (0.58) |
| 500 | 50 | 0.8 | 0.08 | 100 (0.93) | 100 (6.58) |
| 500 | 50 | 1 | 0.08 | 100 (0.94) | 100 (7.74) |
| 500 | 50 | 0.2 | 0.01 | 100 (0.97) | 100 (18.57) |
| 500 | 50 | 0.4 | 0.01 | 100 (0.99) | 100 (47.17) |
| 500 | 50 | 0.8 | 0.01 | 100 (1.0) | 100 (177.0) |
| 500 | 50 | 1 | 0.01 | 100 (1.0) | 100 (324.56) |

**Table S4.** Both sexes are choosy. Mating with replacement. *N* is the population size and *n* the sample size of matings. Choice implies preference for similarity. The quantitative trait distribution is *Z*(0,1). Number of runs: 1,000. % sig: The % of tests with *p*-value ≤0.05. The values in parentheses correspond to the average value of the statistic for the total number of significant runs.

| $N$ | $n$ | Choice | $s$ | % sig. ( $\rho$ ) | %sig. ($\rho^2/(1-\rho^2)$ ) |
| --- | --- | --- | --- | --- | --- |
| $10^4$ | 500 | 0 | -- | 4.4 (0.03) | 5.4 (0.06) |
| $10^4$ | 500 | 0.2 | 0.08 | 24.6 (0.25) | 26.1 (0.07) |
| $10^4$ | 500 | 0.4 | 0.08 | 100 (0.47) | 100 (0.3) |
| $10^4$ | 500 | 0.8 | 0.08 | 100 (0.75) | 100 (1.39) |
| $10^4$ | 500 | 0.2 | 0.01 | 100 (0.93) | 100 (7) |
| $10^4$ | 500 | 0.4 | 0.01 | 100 (0.98) | 100 (26) |
| $10^4$ | 500 | 0.8 | 0.01 | 100 (1) | 100 (106) |
| 500 | 50 | 0 | -- | 4.5 (0.017) | 5.7 (0.12) |
| 500 | 50 | 0.2 | 0.08 | 36.3 (0.36) | 38.5 (0.16) |
| 500 | 50 | 0.4 | 0.08 | 99.8 (0.58) | 99.9 (0.57) |
| 500 | 50 | 0.8 | 0.08 | 100 (0.81) | 100 (2.1) |
| 500 | 50 | 0.2 | 0.01 | 100 (0.96) | 100 (12.7) |
| 500 | 50 | 0.4 | 0.01 | 100 (0.99) | 100 (56.7) |
| 500 | 50 | 0.8 | 0.01 | 100 (1.0) | 100 (206) |

**Table S5.** Comparison of ρ and *J*_*PSI*_ for lower choice values. One sex is choosy. Choice implies preference for similarity. Mating is with replacement. *N* is the population size and *n* the sample size of matings. The quantitative trait distribution is *Z*(0,1). Number of runs: 10,000. % sig: The % of tests with *p*-value ≤0.05. The values in parentheses correspond to the average value of the statistic for the total number of significant runs.

| $N$ | $n$ | Choice | $s$ | % sig. ( $\rho$ ) | %sig. ($\rho^2/(1-\rho^2)$ ) |
| --- | --- | --- | --- | --- | --- |
| 500 | 50 | 0.05 | 0.08 | 5.1 (0.05) | 6.2 (0.12) |
| 500 | 50 | 0.1 | 0.08 | 11 (0.32) | 13 (0.13) |
| 500 | 50 | 0.2 | 0.08 | 38.6 (0.37) | 41.7 (0.16) |
| 500 | 50 | 0.3 | 0.08 | 86 (0.44) | 88 (0.27) |
| 500 | 50 | 0.4 | 0.08 | 99.95 (0.61) | 99.97 (0.64) |
| 500 | 50 | 0.5 | 0.08 | 100 (0.71) | 100 (1.1) |
| 500 | 50 | 0.05 | 0.01 | 99.9 (0.64) | 100 (0.78) |
| 500 | 50 | 0.1 | 0.01 | 100 (0.90) | 100 (4.6) |
| 500 | 50 | 0.2 | 0.01 | 100 (0.96) | 100 (13.54) |
| 500 | 50 | 0.3 | 0.01 | 100 (0.99) | 100 (47.6) |
| 500 | 50 | 0.4 | 0.01 | 100 (0.99) | 100 (56.68) |
| 500 | 50 | 0.5 | 0.01 | 100 (1.0) | 100 (188.6) |

**Figure S1.**
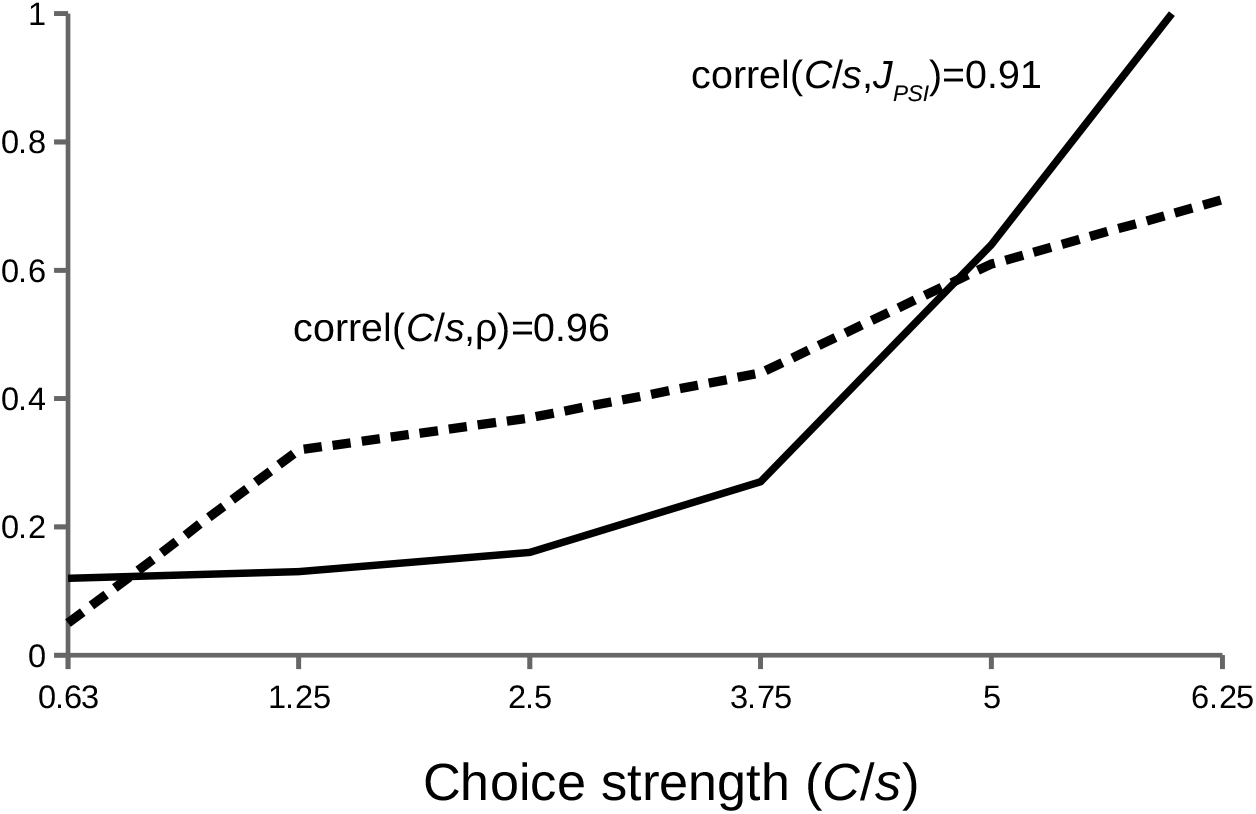
Graphic representation of values of the rho (dashed line) and *J*_*PSI*_ (continuous line) statistics with respect to the strength of the choice. The value of the correlation between each statistic and the strength of the choice is given. Population size *N*=500, sample size *n*=50. The values of ρ and *J*_*PSI*_ are those in Table S5 ordered by increasing *C*/*s*.

